# NfκBin: A Machine Learning based Method for Screening Nf-κB Inhibitors

**DOI:** 10.1101/2025.01.05.630984

**Authors:** Shipra Jain, Ritu Tomer, Sumeet Patiyal, Gajendra P. S. Raghava

## Abstract

Nuclear Factor kappa B (NF-κB) is a transcription factor whose upregulation is associated in chronic inflammatory diseases, including rheumatoid arthritis, inflammatory bowel disease, and asthma. In order to develop therapeutic strategies targeting NF-κB-related diseases, we developed a computational approach to predict drugs capable of inhibiting NF-κB signaling pathways. In this study, we utilized a dataset comprising 1,149 inhibitors and 1,332 non-inhibitors retrieved from PubChem. Chemical descriptors were computed using the PaDEL software, and relevant features were selected using advanced feature selection techniques. Initially, machine learning models were constructed using 2D descriptors, 3D descriptors, and molecular fingerprints, achieving maximum AUC values of 0.66, 0.56, and 0.66, respectively. To improve feature selection, we applied univariate analysis and SVC-L1 regularization to identify features that can effectively differentiate inhibitors from non-inhibitors. Using these selected features, we developed machine learning models, our support vector classifier achieved an AUC of 0.75 on the validation dataset. All models were trained using five-fold cross-validation and rigorously evaluated on an independent validation dataset, which was not used during training or hyperparameter optimization. Finally, our best-performing model was employed to screen FDA-approved drugs for potential NF-κB inhibitors. Notably, most of the predicted inhibitors corresponded to drugs previously identified as inhibitors in experimental studies, underscoring the model’s predictive reliability. Our best-performing models have been integrated into a standalone software and web server, NfκBin, to enable the scientific community to perform high-throughput screening of Nf-κB inhibitors from chemical libraries. (https://webs.iiitd.edu.in/raghava/nfkbin/).

**Highlights:**

1) Inhibitors of Nf-κB can be used to manage disease like rheumatoid arthritis, asthma.
2) A novel tool NfκBin developed for screening Nf-κB inhibitors from chemical library.
3) Cutting-edge feature selection techniques used to identify the most relevant chemical descriptors.
4) Models were developed and validated on a robust dataset of inhibitors and non-inhibitors.
5) Integrated best-performing models into the NfκBin software and web server.

## Introduction

Nuclear factor kappa B (Nf-κB) is a pivotal transcription factor that regulates genes critical for immune and inflammatory responses [1]. Since its discovery in 1986, Nf-κB has been identified as central to the body’s defense mechanisms [2]. It is activated by receptors such as Toll-like receptors (TLRs), which detect microbial components and trigger inflammatory in response to harmful stimuli like pathogens, damaged cells, and irritants [3–5]. Nf-κB activation occurs via two main pathways: canonical and non-canonical [6–10]. The canonical pathway, triggered by signals such as TNF-α and IL-1, involves the phosphorylation and degradation of IκB, allowing Nf-κB to translocate into the nucleus and initiate transcription of genes related to inflammation and immunity [1, 11, 12]. The non-canonical pathway, activated by receptors like CD40 and BAFF, relies on NIK-mediated processing of p100 into its active form (p52), which pairs with RelB to promote immune system development and adaptive immune responses [7].

**Figure 1:**
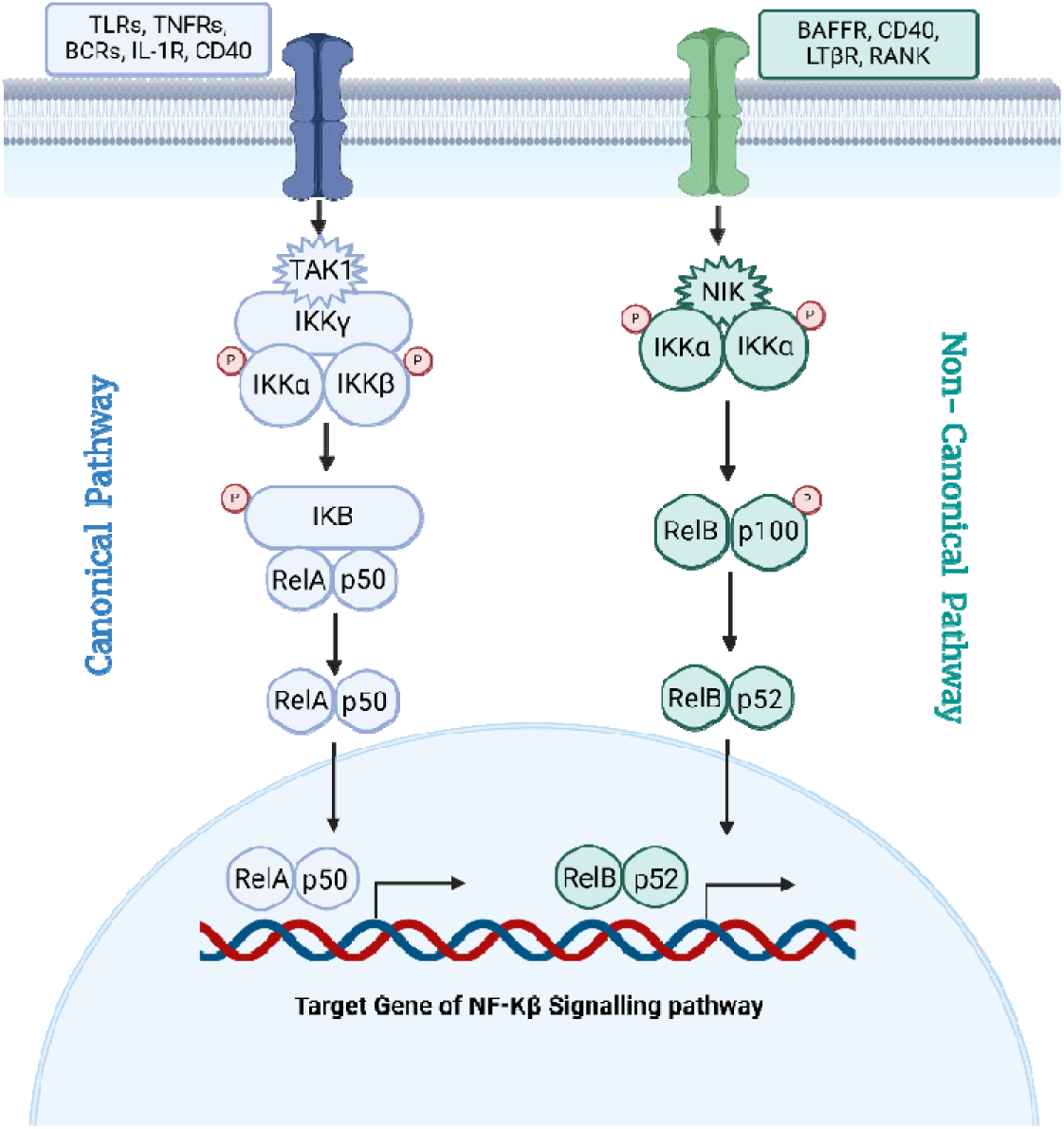
Schematic representation of canonical and non-canonical signaling pathway for activation of Nf-κB.

Dysregulated Nf-κB signaling is implicated in numerous diseases, including chronic inflammatory conditions (e.g., Crohn’s disease, asthma, and psoriasis), autoimmune disorders (e.g., SLE and multiple sclerosis), and cancers (e.g., breast, lung, and colorectal cancers) [9, 13, 14]. Persistent activation contributes to excessive inflammation, tissue damage, and tumor progression by promoting cell survival, proliferation, and resistance to apoptosis [15, 16]. Moreover, it fosters tumor invasion, metastasis, and angiogenesis by creating a pro-tumor microenvironment [17–22].

Due to its central role in diverse diseases, Nf-κB is a promising therapeutic target [8, 23]. Existing inhibitors range from small molecules to natural compounds and peptides, targeting various stages of Nf- κB signaling [24]. However, traditional drug development methods are expensive and time-consuming. Computational approaches for high-throughput screening of chemical libraries to identify Nf-κB inhibitors are urgently needed. In this study, we introduce “NFκBIn,” an in-silico tool for predicting Nf- κB inhibitors based on experimentally validated compounds. This tool addresses the gap in computational resources for efficient and precise inhibitor prediction.

## Materials and Methods

### Dataset Collection

In this study, we extracted the TNF alpha induced Nf-κB inhibitors and non-inhibitors from the PubChem repository [25]. We filtered all the assays in the aforementioned repository using keywords “((TNF AND Nf-κB) inhibitors)”. This search resulted in a total of 90 PubChem bioassays, which was further manually refined based on the number of inhibitors per assay. After rigorous screening we selected a high throughput bioassay AID 1852 (https://pubchem.ncbi.nlm.nih.gov/bioassay/1852) as the data source for our study. This high throughput assay is designed for identification of hits specific to tumor necrosis factor alpha (TNF-a), a canonical Nf- κB inducer, and its modulated pathways. The HEK-293-T Nf-κB-Luc cell line, engineered for luminescent detection of Nf-κB activation, was used in that assay. In this assay, experiment was performed using 0.62% concentration of DMSO supplemented over NOD plating media. Tiered Activity Scoring System developed by SBCCG, was deployed and the compounds showing more than 50% activity in the assay were classified as active. Using this assay, we downloaded a total of 2481 compounds in which 1332 were non-inhibitors and 1149 were reported as Nf-κB inhibitors, using this bioassay.

### Dataset Preprocessing

In this study, we followed the best practices of machine learning algorithms and divided our total compound dataset in 80:20 ratio. Where, 80% of data (i.e. 936 inhibitors and 1048 non-inhibitors) was flagged as training data and were utilized to develop machine learning models and remaining 20% data (i.e. 213 inhibitors and 284 non-inhibitors) was used as independent validation set for machine learning model performance evaluation. These types of standard protocols were reported in previous studies from literature [26–28].

### Molecular Descriptors and Fingerprints of Compounds

Molecular descriptors and fingerprints are the mathematical representation of chemical compounds that captures vital information about them. The descriptors are key features extracted to represent chemical compounds in computational chemistry and drug discovery. They help in predicting the biological activity, physicochemical properties, and toxicity of compounds. In this study, we deployed PaDEL software [29] for calculation of molecular and fingerprint descriptors of Nf-κB inhibitors and non-inhibitors downloaded in SMILES format. This software calculated 1875 descriptors including 1444 1D, 2D; 431 3D and 12 types of fingerprints (total 16092 bits). These 17967 2-D, 3-D, and FP descriptors were further screened to develop machine learning algorithms.

### Descriptor Features Preprocessing

The 17,967 generated descriptors exhibited varying range values. To normalize them, we applied the Standard Scaler from the Scikit-learn package, which operates using the z-score algorithm. Post this step, we discarded the descriptors with more than 80% null values. After this we were left with 1107 2D, 431 3D and 9324 FP descriptors, making a total of 10862 descriptors/ features for the dataset.

### Significant Descriptor Selection and Ranking

In order to develop a robust prediction model with higher accuracy, we need to select the most significant descriptors generated from PaDEL software. Thus, ranking and selecting the significant descriptors from the 10862 descriptors set is an important step. In this study we incorporated two approaches to select and rank relevant descriptors i.e. using correlation analysis and univariate analysis.

**A. Correlation based Descriptor selection:** In this approach, we deployed the Variance Threshold package of Scikit (sklearn.feature_selection) to remove the low-variance features from 10862 descriptor set. After eliminating low variance features, we were left with 6084 descriptors comprising of 786 2D, 169 3D and 5129 FP features (Refer Supplementary Table 1). We applied a correlation-based feature selection method to further screen highly correlated features with a cut off value of 0.6 [30]. Post this step, we were left with 102 2D, 3 3D and 2260 FP descriptors making a total of 2365 descriptors (See Supplementary Table 2). In order to further reduce the dimensionality of the descriptor matrix, we applied SVC-L1 based feature selection method to screen relevant feature set. The support vector classifier (SVC) with linear kernel and L1 regularization is the foundation of this approach [31]. Using SVC-L1 feature selection method we selected, 32 2D, 3 3D and 348 FP feature set (Refer Supplementary Table 3 for list). Using these descriptors, we developed 2D, 3D, FP and ensemble-based machine learning models to screen Nf-κB inhibitors.

**B. Univariate analysis-based Descriptor selection:** In this approach, a statistical method i.e. univariate analysis using 2-tailed independent Student’s t-test was executed based on the mean value of descriptors of both groups to extract the important descriptors from the 10862 descriptors pool. Using this approach, we ranked the descriptors based on the significant p-value obtained. We selected top 2000 descriptors and applied SVC-L1 and RFE based feature extraction methods over them. Recursive Feature Elimination (RFE) is a feature selection approach that works by recursively eliminating the least important features, this process continues until the desired number of features is reached [32]. Applying these, we selected 266 descriptors from SVC-L1 method and the top 50 descriptors from RFE feature selection technique for machine learning model development.

### Cross Validation Techniques

In order to achieve an unbiased prediction model, we incorporated standard five-fold cross validation techniques, to build our machine learning models. In this technique, we divided our 80% training dataset into five sets of data with similar size. Out of these five sets, four sets were used to train the machine learning model and one set was used for testing the machine learning model performance. This process was repeated five times, to make sure that each fold is used once for testing the model. We fine-tuned the machine learning models parameters for achieving best performance on the test dataset. Finally, the average performance was computed using five test folds performance.

### Machine learning models

In this study we have applied various machine learning algorithms to develop prediction models for screening of Nf-κB inhibitors and non-inhibitors with higher accuracy. We implemented Random Forest (RF), Decision Tree (DT), K-nearest neighbour (KNN), Support Vector Classifier (SVC), and eXtreme Gradient Boosting (XGB) to develop classification models. These machine learning algorithms were deployed using the Scikit-learn package [33].

### Performance Evaluation

We have evaluated our machine learning model performance over 20% independent validation dataset. We recorded both threshold-dependent and independent parameters for evaluating our model’s performance. Sensitivity (Sens), specificity (Spec), accuracy (Acc), and Matthew’s correlation coefficient (MCC) were recorded as threshold-dependent parameters and the area under the receiver operating characteristic curve (AUC), as the threshold-independent parameter [34,35].

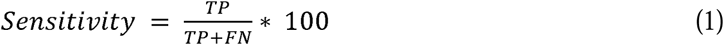

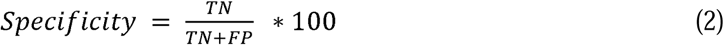

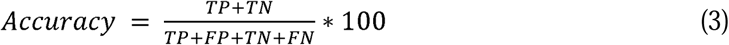

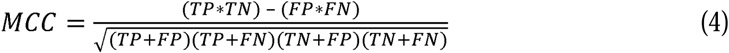

Where, FP is false positive, FN is false negative, TP is true positive and TN is true negative.

## Results

### Functional Group analysis

In order to get the deeper insights of relevance of functional groups present in Nf-κB signaling pathway inhibitors and non-inhibitors. We used ChemmineR package to detect and the frequency of functional groups in both Nf-κB inhibitors and non-inhibitors chemical compounds [36]. Using this approach, we observed the occurrence of Primary (RNH2), Secondary (R2NH), Tertiary (R3N) Amines, Phosphates attached to alkyl groups (ROPO3), Alcohol (ROH), Aldehyde (RCHO), Ketone (RCOR), Carboxylic Acid (RCOOH), Ester (RCOOR), Ether (ROR), Alkyne (RCCH), Nitrile (RCN), Rings and Aromatic groups for our positive and negative dataset. The frequency of these functional groups is depicted in Fig 2.

**Figure 2:**
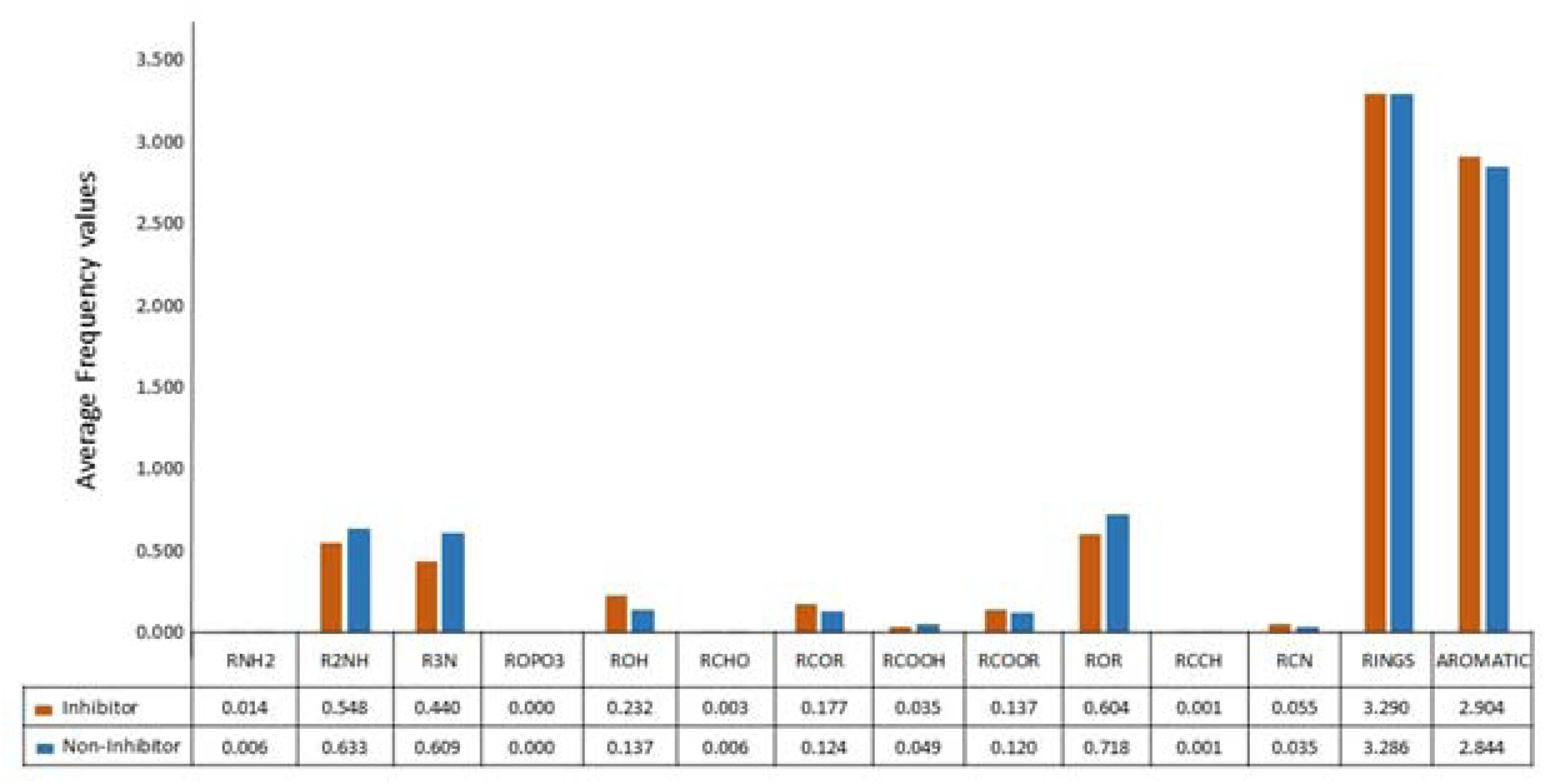
Representation of average functional groups frequency reported in Nf-κB pathway inhibitors and non-inhibitors using ChemmineR package.

We observed that the alcohol (ROH) functional group had a higher average frequency value in inhibitor compounds as compared to non-inhibitors. Whereas, Secondary amines (R2NH), Tertiary amines (R3N) and Ether (ROR) functional groups was found in abundance in non-inhibitor molecules.

### Performance of prediction models

**A. Correlation based descriptor model performance:** In this approach, after eliminating highly correlated features, we selected 32 2D, 3 3D and 348 FP descriptors using SVC-L1 based feature selection method. We developed 2D, 3D, FP and ensemble-based machine learning prediction model.

I. **2D descriptors-based ML model:** Machine learning prediction model was developed using 32 2D descriptors. Using this approach, K-Nearest Neighbor method recorded the maximum AUC for validation set as 0.62 and accuracy as 64.85%. Performance of 2D descriptors over training dataset can be referred in Supplementary Table 4.

**Table 1:** The machine-learning model performance on validation dataset developed using 32 2D descriptors.

|  | Validation |  |  |  |  |  |  |  |
| --- | --- | --- | --- | --- | --- | --- | --- | --- |
| Model | Accuracy | Precision | Recall | F1 | Sens | Spec | AUC | MCC |
| RF | 65.05 | 0.65 | 0.49 | 0.74 | 0.49 | 0.56 | 0.61 | 0.24 |
| DT | 54.23 | 0.54 | 0.44 | 0.63 | 0.44 | 0.49 | 0.54 | 0.07 |
| KNN | 64.85 | 0.65 | 0.53 | 0.72 | 0.53 | 0.58 | 0.62 | 0.25 |
| SVC | 61.33 | 0.61 | 0.37 | 0.77 | 0.37 | 0.46 | 0.57 | 0.15 |
| XGB | 61.33 | 0.61 | 0.45 | 0.72 | 0.45 | 0.52 | 0.58 | 0.18 |

II. **3D descriptors-based ML model:** We have also developed machine learning based model using 3 3D descriptors screened. We observed the maximum AUC over validation set as 0.56 with accuracy as 56.14% in Random Forest classifier. Performance of 3D descriptors over training dataset can be referred in Supplementary Table 4.

**Table 2:** The machine-learning model performance on validation dataset developed using 3 3D descriptors.

|  | Validation |  |  |  |  |  |  |  |
| --- | --- | --- | --- | --- | --- | --- | --- | --- |
| Model | Accuracy | Precision | Recall | F1 | Sens | Spec | AUC | MCC |
| RF | 56.14 | 0.57 | 0.49 | 0.64 | 0.49 | 0.52 | 0.56 | 0.12 |
| DT | 52.31 | 0.52 | 0.55 | 0.50 | 0.55 | 0.53 | 0.52 | 0.05 |
| KNN | 53.52 | 0.53 | 0.55 | 0.52 | 0.55 | 0.54 | 0.54 | 0.07 |
| SVC | 51.51 | 0.52 | 0.32 | 0.71 | 0.32 | 0.40 | 0.51 | 0.03 |
| XGB | 51.91 | 0.68 | 0.06 | 0.97 | 0.06 | 0.11 | 0.52 | 0.08 |

III. **Fingerprint descriptors-based ML model:** In this study, we developed classification models using 348 fingerprints descriptors selected using correlation and SVC-L1 based feature selection approach. Using FP descriptors, we achieved a maximum AUC of 0.66 and accuracy as 66.40% over validation dataset using Random Forest classifier. Also, XGBoost model reported the AUC as 0.66, and 65.79% as accuracy for validation set. Performance of FP descriptors over training dataset can be referred in Supplementary Table 4.

**Table 3:** The machine-learning model performance on validation dataset developed using 348 FP descriptors.

|  | Validation |  |  |  |  |  |  |  |
| --- | --- | --- | --- | --- | --- | --- | --- | --- |
| Model | Accuracy | Precision | Recall | F1 | Sens | Spec | AUC | MCC |
| RF | 66.40 | 0.74 | 0.51 | 0.82 | 0.51 | 0.60 | 0.66 | 0.34 |
| DT | 57.95 | 0.59 | 0.50 | 0.66 | 0.50 | 0.54 | 0.58 | 0.16 |
| KNN | 65.19 | 0.66 | 0.62 | 0.69 | 0.62 | 0.64 | 0.65 | 0.30 |
| SVC | 59.96 | 0.61 | 0.54 | 0.66 | 0.54 | 0.57 | 0.60 | 0.20 |
| XGB | 65.79 | 0.68 | 0.59 | 0.73 | 0.59 | 0.63 | 0.66 | 0.32 |

IV. **Ensemble based approach:** In order to improve the machine learning model performance, we adopted an ensemble-based approach in this study. We combined 32 2D, 3 3D and 348 FP descriptor set selected using SVC-L1 feature selection approach applied after removing highly correlated features. We developed a machine learning model, using a matrix of 383 feature set of 2D, 3D, FP descriptors. We recorded the maximum AUC as 0.67 and accuracy of 67.20% using Random Forest classifier over validation dataset. Performance of FP descriptors over training dataset can be referred in Supplementary Table 4.

**Table 4:** The machine-learning model performance on validation dataset developed using 383 ensemble-based descriptors set.

|  | Validation |  |  |  |  |  |  |  |
| --- | --- | --- | --- | --- | --- | --- | --- | --- |
| Model | Accuracy | Precision | Recall | F1 | Sens | Spec | AUC | MCC |
| RF | 67.20 | 0.73 | 0.53 | 0.81 | 0.53 | 0.62 | 0.67 | 0.36 |
| DT | 56.54 | 0.57 | 0.51 | 0.62 | 0.51 | 0.54 | 0.57 | 0.13 |
| KNN | 64.79 | 0.66 | 0.62 | 0.68 | 0.62 | 0.63 | 0.65 | 0.30 |
| SVC | 60.97 | 0.62 | 0.54 | 0.68 | 0.54 | 0.58 | 0.61 | 0.22 |
| XGB | 64.59 | 0.68 | 0.54 | 0.75 | 0.54 | 0.60 | 0.65 | 0.30 |

**B. Univariate analysis-based descriptor model performance:** In this approach, we screened top 2000 descriptors using Univariate analysis. We calculated the mean difference of descriptor score and the single descriptor-based AUC score for top 20 descriptors, refer to Supplementary Table 7. In this, we observed the KRFP605 outperformed all and have shown the maximum AUC as 0.62, with average mean difference as 2.60 among positive and negative data descriptor. In addition to this, we applied SVC-L1 and RFE based feature selection technique over top 2000 descriptors screened using Univariate analysis. We developed machine learning based model for prediction of Nf-κB inhibitors using 266 descriptors from SVC-L1 method and the top 50 descriptors from RFE feature selection technique (Refer Supplementary Table 5 for list). As depicted in Table 5, Support vector classifier (SVC) developed using SVC-L1 based feature selection technique outperformed all classifiers and reported maximum AUC of 0.80 on training dataset and 0.75 on validation dataset. However, K-nearest neighbor classifier reported maximum AUC of 0.66 on training dataset and 0.65 on testing dataset developed using 50 RFE selected descriptors (See Supplementary Table 6).

**Table 5:** The machine-learning models performance on validation dataset developed using 266 descriptors selected using SVC-L1 based approach.

| Model | Threshold | Validation |  |  |  |  |  |  |  |
| --- | --- | --- | --- | --- | --- | --- | --- | --- | --- |
|  |  | Acc | FPR | Kappa | F1 | Sens | Spec | AUC | MCC |
| RF | 0.47 | 65.59 | 29.60 | 0.31 | 0.64 | 60.73 | 70.40 | 0.71 | 0.31 |
| DT | 0.48 | 60.36 | 33.60 | 0.21 | 0.58 | 54.25 | 66.40 | 0.63 | 0.21 |
| KNN | 0.36 | 65.19 | 30.40 | 0.30 | 0.63 | 60.73 | 69.60 | 0.70 | 0.31 |
| SVC | 0.41 | 67.61 | 28.00 | 0.35 | 0.66 | 63.16 | 72.00 | 0.75 | 0.35 |
| XGB | 0.43 | 62.78 | 34.00 | 0.26 | 0.61 | 59.51 | 66.00 | 0.69 | 0.26 |

### FDA approved drug repurposing to target Nf-κB signaling pathway

In this study, we attempted a systemic approach to identify the potential drug targets for Nf-κB signaling pathway. In order to achieve this, we retrieved the 2616 FDA approved drug molecules from Drug Bank portal to screen them as the Nf-κB pathway inhibitors and non-inhibitors. Out of 2616, SMILES format was available for 2577 drug molecules. We deployed the “Predict” module of our NfκBIn webserver over these 2577 compounds SMILES format dataset using the the default parameters. It computed descriptors using PaDEL software in the backend and provided a machine learning based model label for each drug candidate in 2577 compounds dataset. The machine learning score and predicted label as inhibitor or non-inhibitor for 2577 compounds can be referred in Supplementary Table 8. For top 10 potential drug candidates identified as Nf-κB signaling pathway inhibitors, we reviewed previous studies to validate and support our findings. These studies provide evidence of the inhibitory effects of six compounds on the Nf- κB pathway, reinforcing the potential of these candidates for further investigation and development [37–43]. These seven potential drugs i.e. Bleomycin, Nitroprusside, Guanidine Ivermectin, Tobramycin, Pentosan polysulfate and Gentamicin and their roles as reported in various studies are depicted in Table 6.

**Table 6:**
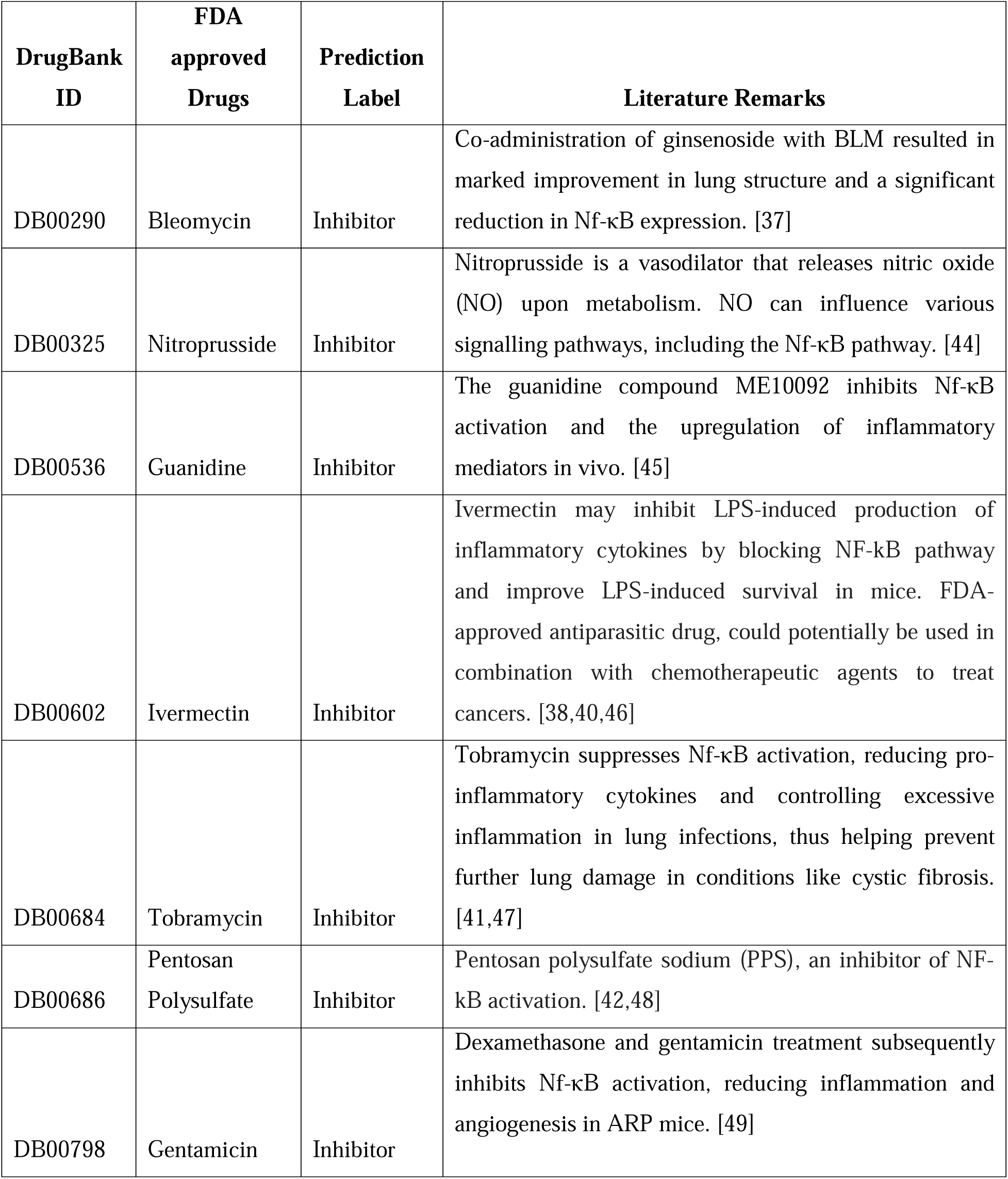
List of seven FDA-approved drug candidates as potential Nf-κB pathway inhibitors.

### Webserver and Standalone Package

In this study, we have provided a user-friendly webserver “NFκBin” (https://webs.iiitd.edu.in/raghava/nfkbin/)platform to enable high-throughput screen of chemical compounds as Nf-κB inhibitors and non-inhibitors. This webserver is deployed on a Linux (Ubuntu) machine using an Apache HTTP server. Its front-end is created with HTML, PHP, and JavaScript, while the back-end is implemented in Python 3.6 utilizing the Scikit library. In addition to this, to ease the usability of the webserver we have utilized a responsive template which is compatible with desktop, tablet and phone. Major modules incorporated in this webserver, are “Predict,” “Draw,” and “Analog design”. Predict module enables users to screen the chemical compounds in SMILES format as Nf-κB inhibitors and non-inhibitors. Best machine learning model has been incorporated in this module with default threshold parameter. Draw module allows users to draw or modify the chemical compound’s structure using an open-source interactive tool known as Ketcher. Post that, drawn structure can be further classified as Nf-κB inhibitors and non-inhibitors. In order to generate the analog’s of the chemical compounds using combination of scaffolds, building blocks, and linkers, users can utilize the Analog Design module. SmiLib tool has been implemented in the backend of this module. The tabular format results generated can be downloaded in .csv format from all modules. In addition, we also developed standalone software package which is available from GitHub and PyPI site (https://github.com/raghavagps/nfkbin/ & https://pypi.org/project/nfkbin/).

**Figure 3:**
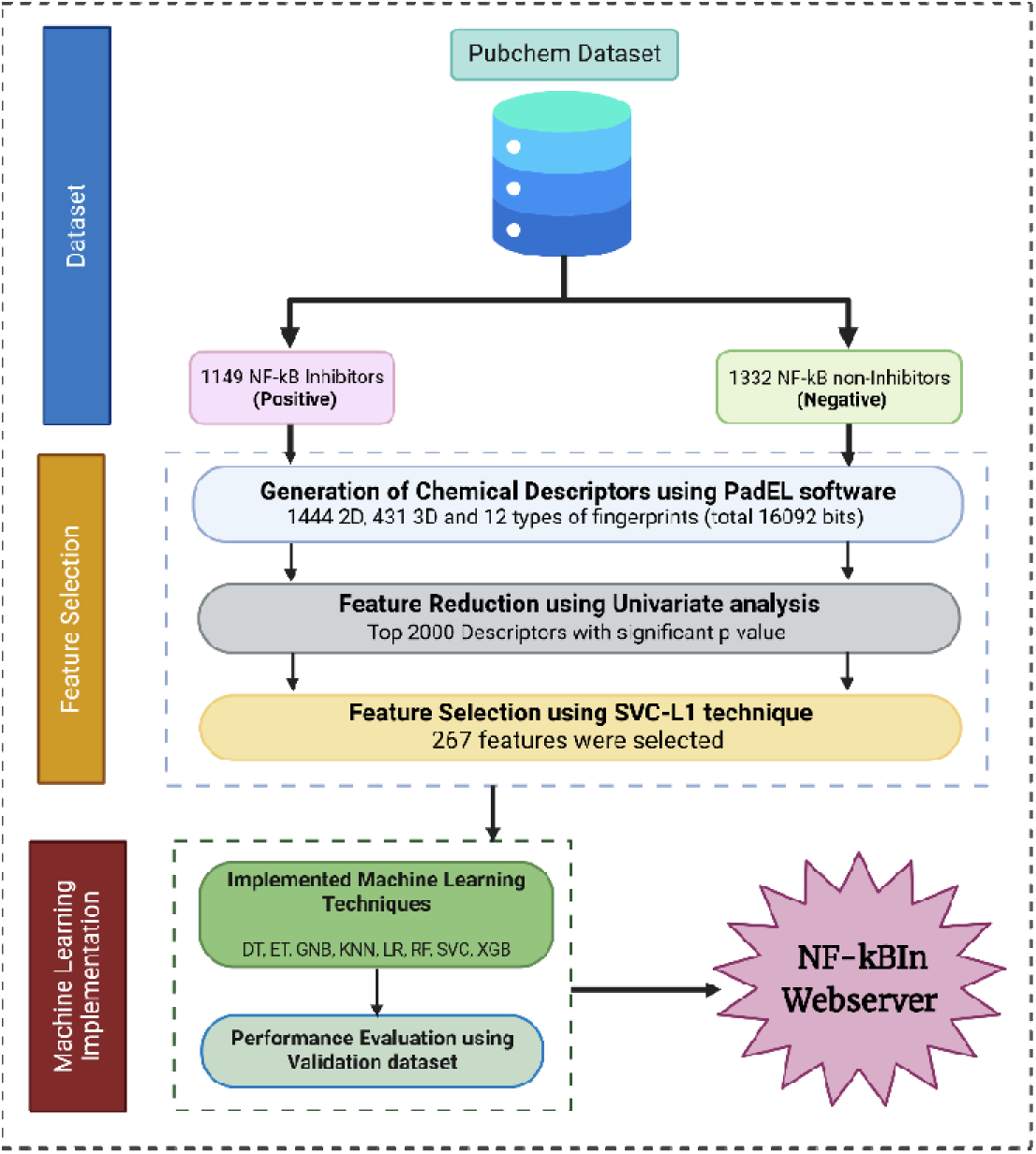
Overall architecture depicting workflow of Nf-κBIn tool.

## Discussion

Nf-κB is a vital therapeutic target in drug discovery fields, due to its abnormal activation reported in several diseases particularly related to chronic inflammation, immune system dysfunction, and uncontrolled cell growth. TNF alpha activates Nf-κB molecules via canonical pathway, which leads to its translocation into the nuclear membrane and further it enables transcription of genes. Researchers have emphasized on the need of blocking Nf-κB signaling pathway at initial stages, so that its abnormal upregulation in diseases can be modulated. Several tools developed in past focusing on various broader domain such as EGFRpred [50] aims to predict the potential chemical molecule as an EGFR inhibitor based on the structure-activity (QSAR model) of the chemical compound; DrugMint [51] to scan and identify whether a chemical molecule is a potential drug candidate or not; ChAlPred [28] tool for predicting allergenicity of chemical compounds.

In addition to these, several molecular docking and simulation-based studies have been conducted for screening of chemical compounds as Nf-kB inhibitors [52–59]. Traditional molecular docking methods are often limited due to their focus on individual receptor targets, overlooking the broader context of pathway-level interactions. Additionally, they have limited accuracy due to dynamic nature of protein-ligand interactions and the vastness of the chemical space, leading to high false-positive rates and reduced predictive accuracy. In contrast, machine learning-based approaches target entire pathways, leveraging larger datasets with physicochemical descriptor information, leading to improved efficiency and accuracy of identifying potential Nf-κB inhibitors.

In present study, we presented “NFκBIn”, an in-silico computational tool for high throughput screening of chemical compounds libraries as the Nf-κB inhibitors and non- inhibitors with accuracy. This computer aided tool is built using a machine learning algorithm, analyzing a molecular descriptors and fingerprints of 2481 experimentally validated compounds. Out of these 2481 compounds, 1332 were non-inhibitors, while 1149 were flagged as specific inhibitors of TNF-α-mediated Nf-κB activation. Moving forward, we hope that this state-of-the-art tool will be used by the researchers working in computational drug discovery field for identification of novel drugs against disease caused by TNF-α-mediated Nf-κB signaling pathway.

## Abbreviations

Nf-κB: Nuclear factor kappa B
TNF: Tumor Necrosis Factor
FP: Fingerprint descriptor
RFE: Recursive Feature Elimination
MCC: Mathews Correlation Coefficient
DT: Decision Tree
RF: Random Forest
LR: Logistic Regression
XGB: Extreme Gradient Boosting
KNN: K-nearest neighbour
SVC: Support Vector Classifier
Sens: Sensitivity
Spec: Specificity
Acc: Accuracy

## Future Scope

Nf-κBIn method can be implied in Computational drug discovery pipelines to conduct virtual screening of chemical compound libraries as Nf-κB inhibitors. Repurposing of FDA-approved drugs as potential candidates against the Nf-κB pathway opens new avenues for therapeutic interventions. These findings strengthen the case for further exploration and development of six compounds as viable drug candidates. In addition to this, webserver enable scientific community to create or modify chemical compounds for the discovery of novel chemical compounds targeting against Nf-κB signaling pathway.

## Author Contributions

SJ conceived the idea, collected and processed the datasets. SJ and RT implemented the algorithms and developed the prediction models. SJ, RT and GPSR analyzed the results. SJ and RT created the web server. SP created the standalone and PyPi version of the tool. SJ and GPSR penned the manuscript. GPSR coordinated the project. All authors have read and approved the final manuscript.

## Acknowledgement

Authors are thankful to the Department of Bio-Technology (DBT) for the financial support, and the Department of Computational Biology, IIITD New Delhi for infrastructure and facilities. We would like to acknowledge that Figures were created using BioRender.com.

## Data Availability Statement

All the datasets generated for this study are available at the “NFκBIn” web server, https://webs.iiitd.edu.in/raghava/nfkbin/dataset.php.

